# Neuromorphological changes following selection for tameness and aggression in the Russian fox-farm experiment

**DOI:** 10.1101/2020.11.21.390229

**Authors:** Erin E. Hecht, Anna Kukekova, David A. Gutman, Greg Acland, Todd M. Preuss, Lyudmila Trut

## Abstract

The Russian fox-farm experiment is an unusually long-running and well-controlled study designed to replicate wolf-to-dog domestication. As such, it offers an unprecedented window onto the neural mechanisms governing the evolution of behavior. Here we report adaptations to gray matter morphology resulting from selection for tameness vs. aggressive response toward humans. Contrasting with prior work in other domesticated species, tame foxes did not show reduced brain volume. Rather, gray matter volume in both the tame and aggressive strains was increased relative to foxes bred without selection on behavior. Furthermore, tame- and aggressive-enlarged regions overlapped substantially, including portions of motor, somatosensory, and prefrontal cortex, amygdala, hippocampus, and cerebellum. We also observed differential morphological covariation across distributed gray matter networks. In one prefrontal-hypothalamic network, this covariation differentiated the tame and aggressive foxes together from the conventional strain. These findings indicate that selection for opposite behaviors can influence brain morphology in a similar way.

## Introduction

Domestication refers to the process of animal adaptation to the human niche. It represents one of the largest and most rapid evolutionary shifts in life on Earth: the biomass of domesticated animals has increased an estimated 3.5-fold in the last 100 years and now outweighs the biomass of other terrestrial mammals by a factor of about 25^1^. Correspondingly, the neural changes associated with domestication constitute a major event in the history of brain evolution. Moreover, self-domestication is hypothesized to have played a role in the evolution of our own species^2–4^. However, surprisingly little is known about the neural correlates of domestication.

Perhaps the most well-known effect of domestication on the brain is a reduction in its size^5^. Dogs are the oldest and perhaps the archetypal domesticate, having split from wolves an estimated 10,000-30,000 years ago^6,7^. Past research on the neural correlates of wolf-to-dog domestication has implicated prefrontal cortex and the limbic system, particularly the HPA axis. For example, gene expression in the hypothalamus is conserved between wolves and coyotes but diverges significantly in dogs^8^. Similarly, genes showing high differentiation between wolves and Chinese native dogs show high expression bias for the brain, particularly those expressed in prefrontal cortex^9^. In an MRI study of 8 wild carnivore species and 13 domestic dogs, the allometric scaling of corpus callosum size to total brain size was constant across species, except in the rostral component, which interconnects prefrontal cortices^10^. Moreover, the enlargement of one component of carnivore prefrontal cortex, the prorean gyrus, which has extensive connections with other components of the limbic system^11,12^, has been implicated in the emergence of complex social behavior in canid evolution^13^.

The domestication of wolves into dogs is paralleled by the well-known and long-running Russian fox experiment^14^. Since 1959, researchers at the Institute of Cytology and Genetics in Novosibirsk have been breeding farm-raised foxes on the basis of their behavioral response to human social contact. The tame strain is selected for high social approach behavior toward humans and produces dog-like behaviors such as licking and tail wagging. The aggressive strain is selected for the opposite behavior and reacts with defensive aggression when faced with human contact. A third conventional strain is kept on the farm but bred without selection on behavior^15^. Behavioral differences between tame, aggressive, and conventional foxes map to a locus on fox chromosome 12 which is homologous to the locus implicated in wolf-to-dog domestication^16,17^ and the whole-genome sequencing of the three populations identified 46 genomic regions which are syntenic to canine candidate domestication regions^18^. Differential gene expression across strains has been established in prefrontal cortex, basal forebrain, hypothalamus, and anterior pituitary^19–22^. Additionally, tame foxes show increased adult neurogenesis in the hippocampus^23^. However, the brain-wide neuroanatomical consequences of selection on behavior in the fox model are as yet unknown. In this study, we addressed this question using high-resolution, T2-weighted, whole-brain ex vivo neuroimaging in 10 tame, 10 aggressive, and 10 conventional foxes. These were the same individuals used in previous transcriptomic studies^16,17^, and their behavior was tested and analyzed as previously described.^24^ We carried out three types of analyses: a comparison of overall gray matter, white matter, and total brain volumes; a comparison of differences in regional gray matter volumes across strains; and a comparison of strain-wise differences in anatomical covariation across regions, which can identify morphologically co-evolving structural networks and link them to individual variation in behavior.

## Results

The T2-weighted conventional fox brain template and labels for some anatomical regions are shown in **Figure 1A**. A map of variation in brain anatomy across all strains is shown in **Figure 1B**; a graph of gray matter, white matter, and total brain volumes for each strain are shown in **Figure 1C**. One-way ANOVAs revealed a strain-wise difference in total gray matter volume (F(2,27) = 6.855, p = .004). Post-hoc Tukey’s HSD tests indicated that both tame and aggressive foxes had significantly higher gray matter volume than conventional foxes (tame > conventional: t(18)=3.370, p = .003; aggressive > conventional: t(18)=3.556, p = .002), but did not differ significantly from each other (t(18) = .010, p = .992). Strainwise differences in white matter volume and total brain volume did not reach significance (white matter: F(2,27) = 2.186, p = .132; total brain volume: F(2,27) = 1.504, p = .240).

**Figure 1:**
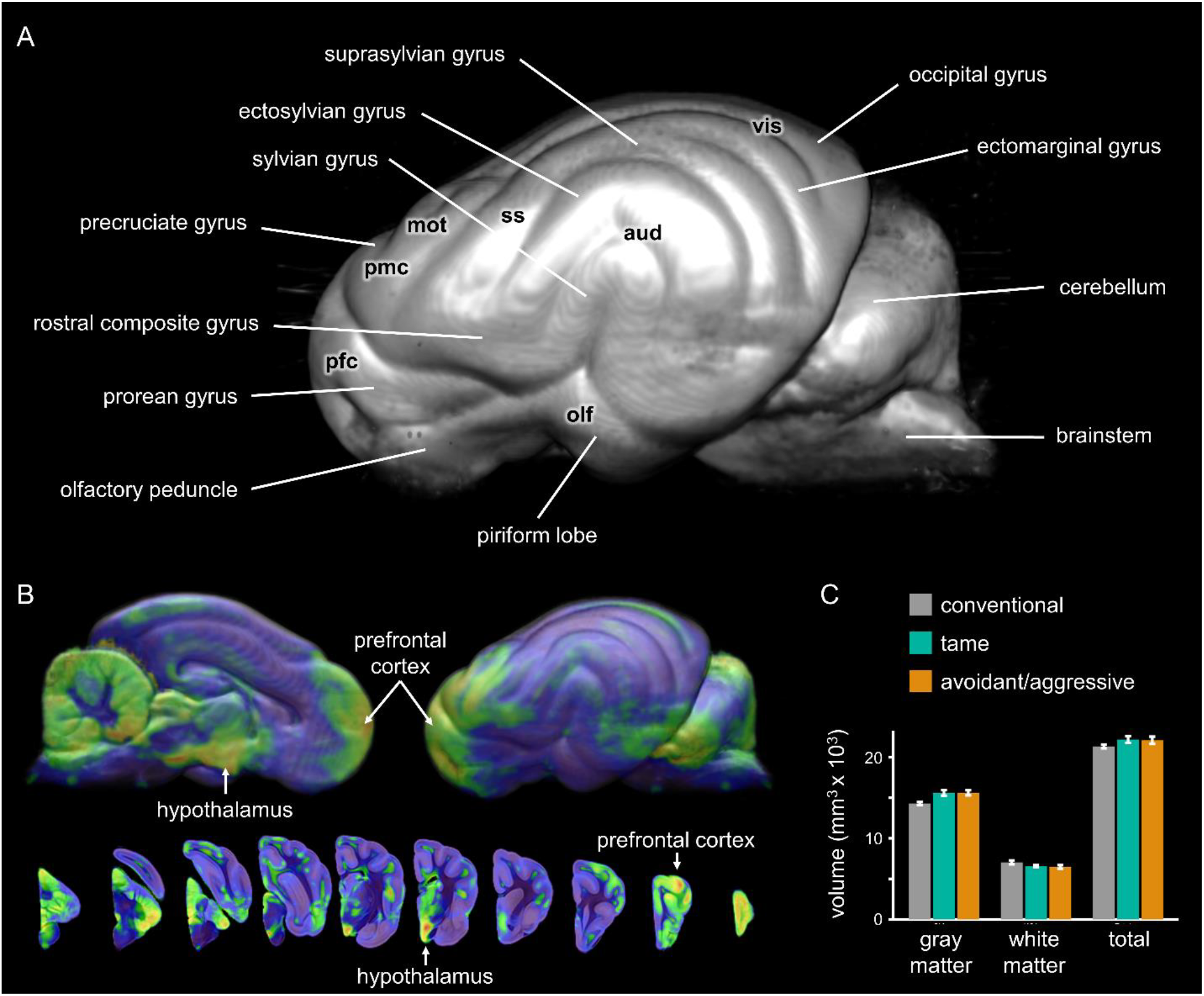
Brain volume shifts in the Russian fox-farm experiment. (A) Group-average template with anatomical labels and approximate cortical functional localizations from the dog^25^. olf: olfactory; pfc: prefrontal; pmc: premotor; mot: motor; ss: somatosensory; aud: auditory; vis: visual. (B) Variation in regional morphology across the dataset (warmer colors indicate more variation). Lateral views of 3D surface renderings and coronal cross-sections are shown. (C) Strain-wise differences in gray matter, white matter, and total brain volume. Error bars: +/− 1 SEM.

Voxel-based morphometry identified a number of differences between strains (**Figure 2**). Carnivore cortical mapping studies have been most extensive in cats and ferrets, but relatively less work has been done in dogs, which are more closely related to foxes. Thus, putative functions of anatomical regions are based on known dog anatomy here, but should be considered tentative. Relative to aggressive foxes, tame foxes show expansion in portions of the sylvian gyrus and ectosylvian sulcus (temporal regions that include auditory cortex and other regions, potentially higher-order visual or multisensory cortex^26^; **Figure 2A**). In contrast, compared to tame foxes, aggressive foxes show expansion in portions of the rostral composite gyrus and presylvian sulcus (potentially somatosensory and/or premotor-prefrontal transition cortex^27^) and in the ectosylvian and sylvian gyri and sylvian sulcus (auditory and association cortex in dogs^26^; **Figure 2B**). Surprisingly, relative to the conventional strain, both tame and aggressive foxes show expansion in similar regions, including portions of the prorean, orbital, frontal, precruciate, and rostral composite gyri (prefrontal, premotor, and motor cortex in dogs^27^), amygdala, hippocampus, and cerebellum (**Figures 2C** and **Figure 2D**; overlap shown in **Figure 2E**). No voxels showed reduced volume in tame or aggressive foxes relative to conventional foxes. Volumes and voxel coordinates for each cluster are shown in **Table S2**.

**Figure 2:**
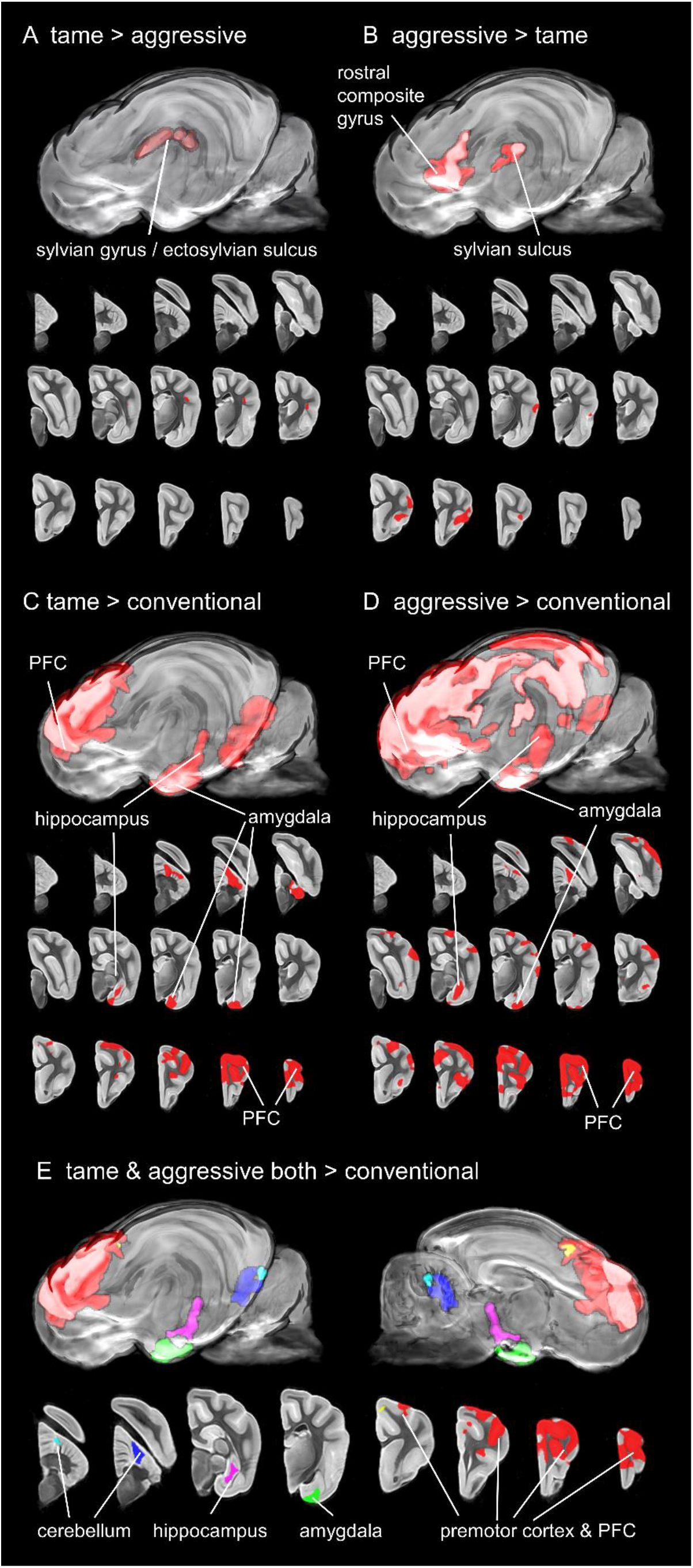
Differences in regional gray matter volume between strains. (A) Tame > aggressive. (B) Aggressive > tame. (C) Tame > conventional. (D) Aggressive > conventional. (E) Tame and aggressive both > conventional. Red: frontal cortex; green: amygdala; magenta: hippocampus; blue and cyan: cerebellum.

Notably, the hypothalamus was one of the regions with the highest volumetric variation across the entire dataset (**Figure 1B**), but direct group-wise comparisons did not identify significant volumetric differences between strains (**Figure 2**). Past studies on the fox-farm experiment have implicated the HPA axis generally (reviewed in^14^) and gene expression in the hypothalamus specifically^21^. Neuroanatomical consequences of rapid selection on behavior can be visible not only in changes to relative volume of brain regions, but also in the degree of morphological correlation across brain regions, as we recently documented in domestic dog breeds^28^. To probe this possibility, we used source-based morphometry, a model-free, ICA-based approach^29^, to identify structurally covarying multi-region networks across the entire dataset (but note that white matter connectivity is not assessed with this method). This identified 4 significantly covarying brain networks (**Figure 3**, **Table S3**).

**Figure 3:**
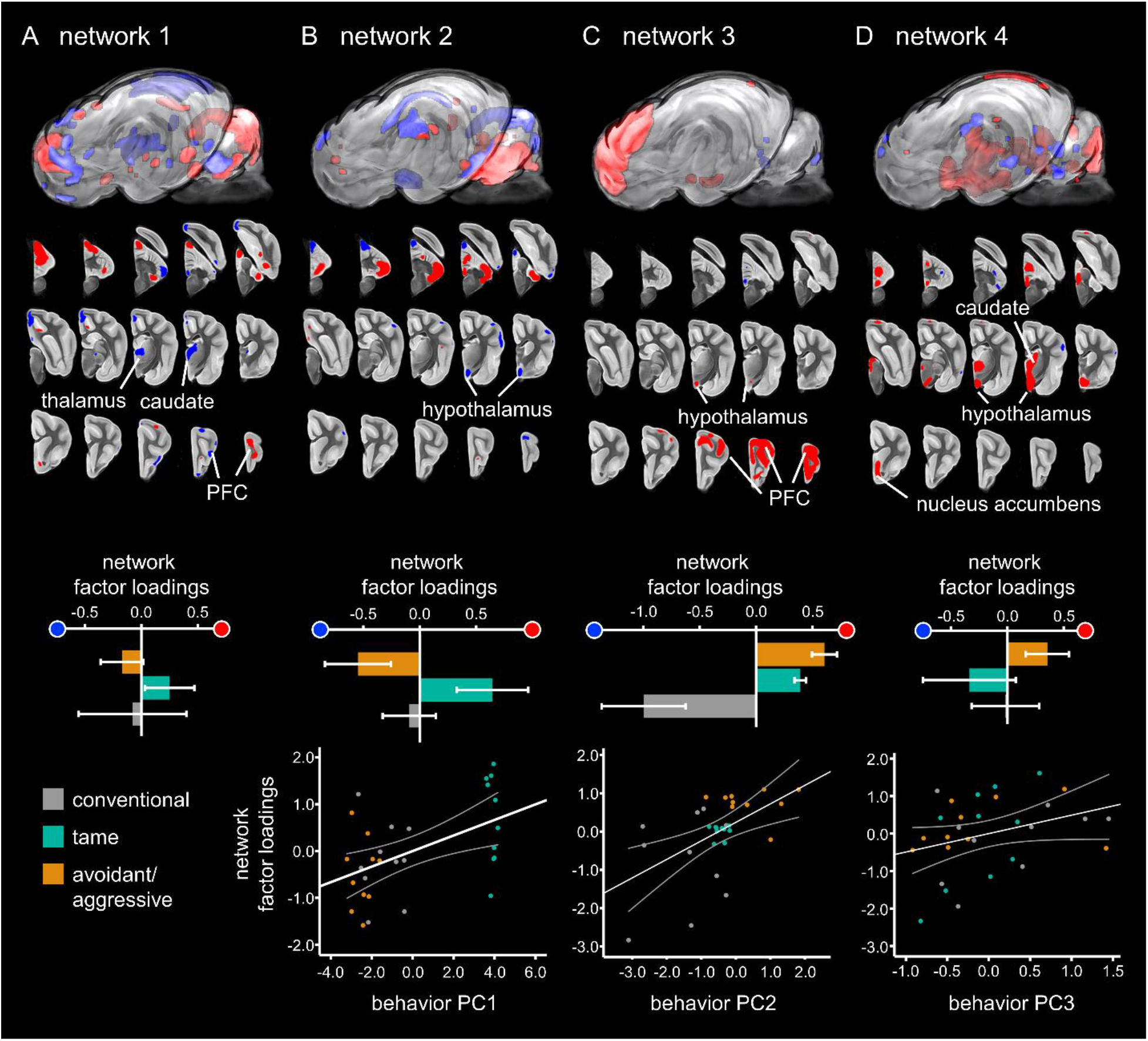
Regionally covarying structural brain networks and relationship to strain membership and individual behavior scores. Red and blue components of each network are anti-correlated. Bar graphs illustrate factor loadings for each strain; positive loadings indicate larger volumes in red regions (indicated with red dot), while negative loadings indicate larger volumes for blue regions (indicated with blue dot). Partial regression plots illustrate significant relationships between factor loadings for networks 2-4 and behavior measurements PC1-3 in each fox. (A) Network 1 factor loadings did not differentiate between strains and were not significantly related to behavior scores. (B) Network 2 factor loadings differentiated between tame and aggressive strains and were significantly related to PC1 behavior scores. (C) Network 3 factor loadings differentiated the tame and aggressive strains from the conventional strain and were significantly related to PC2 behavior scores. (D) Network 4 factor loadings did not differentiate between strains and were marginally related to PC3 behavior scores.

We then investigated the extent to which these networks were related to strain membership and to individual behavior. These were then subjected to a PCA analysis, resulting in 3 components^16^, each of which explains 29.0%, 8.0%, and 6.6% of variance, respectively. We then used multiple linear regression to probe the relationship between factor loadings for Kukekova et al.’s 3 behavioral PCA components (**Table S1**) and our 4 morphometry components (**Table S3**).

Network 1 contained clusters in the thalamus, caudate, nucleus accumbens, cerebellum, and other regions (**Figure 3A**). Factor loadings were in opposite directions for the tame and aggressive strains but were centered near zero with wide variance in the conventional strain; however, strainwise differences did not reach significance (F(2,27) = 0.475, p = 0.627). Factor loadings for Network 1 accounted for 11.6% of the variance in behavior PCA scores, but none of the partial correlations between brain and behavior factor loadings reached significance.

Network 2 was mainly comprised of one cluster that covered most of the hypothalamus, plus several other clusters scattered throughout the cerebellum. Factor loadings strongly differentiated the tame and aggressive strains from each other and were centered near zero for the conventional strain. Strainwise differences were significant (F(2,27) = 4.495, p = 0.012; **Figure 3B**). Post-hoc T-tests confirmed that factor loadings significantly differentiated the tame and aggressive strains from each other (t = −2.764, p = .013). Loading coefficients for network 2 accounted for 31.0% of the variance in behavior scores. There was a significant partial correlation with PC1 behavior scores (β = .166, t = 2.964, p = .006). Generally, behavioral traits with the highest positive loadings for PC1 describe tame behavior and proximity to a human approacher, while traits with the most negative loadings describe aggressive behavior and greater distance from a human approacher (^24^; see **Table S1**). Examination of the partial correlation scatter plot reveals that tame foxes cluster together with higher scores for both behavior and neuroanatomy, while aggressive and conventional foxes cluster together on the lower end of both axes (**Figure 3B**).

Network 3 contained two smaller, discrete clusters in the hypothalamus, plus a large cluster covering much of prefrontal and premotor cortex, including portions of the prorean, orbital, frontal, precruciate, and rostral composite gyri. For this network, factor loadings were similar for the tame and aggressive strains, and both were strongly opposite to the conventional foxes; the strainwise difference was significant (F(2,27) = 14.795, p < .001; **Figure 3C**). This network significantly differentiated both the tame strain from the conventional strain (t = 3.695, p = .002) and the aggressive strain from the conventional strain (t = 4.127, p = .001). Regression analyses revealed that behavior scores accounted for 35.3% of the variance in morphometry loading coefficients. There was a significant partial correlation with PC2 (β = .483, t = 3.251, p = .003). For PC2, traits with high negative loadings describe a passive tame response and tolerance of human tactile contact and fox position in the back of the cage, while traits with high positive loadings describe active aggressive response (^24^; **Table S1**). Examination of the partial correlation scatter plot reveals that aggressive foxes tended to score high for both the behavior and neuroanatomical measures, while conventional foxes tended to score low on both, and tame foxes were tightly clustered in the intermediate range (**Figure 3C**).

Network 4 contained a large cluster that spanned regions of the thalamus, most of the hypothalamus, and portions of the nucleus accumbens/ventral forebrain and caudate (**Figure 3D**). There were also several additional clusters located in cortex and cerebellum. Factor loadings were again opposite for the tame and aggressive strains, and centered near zero for the conventional strain; strainwise differences did not reach significance (F(2,27) = 1.198, p = 0.312). Loading coefficients for Network 4 accounted for 14.6% of the variance in behavior scores. None of the partial correlations between morphometry and behavior scores reached significance, but there was a marginally positive relationship with PC3 (β = .492, t = 1.780, p = .087). In PC3, traits with the highest positive loadings describe neutral exploratory behavior (e.g., “ears are vertical”), while traits with the most negative loadings describe prosocial greeting behavior without tactile contact and fox position mostly close to the experimenter (^24^; see **Table S1**). Tame, aggressive, and conventional foxes showed marked overlap in both behavior and neural scores in this analysis (**Figure 3D**).

## Discussion

The Russian fox-farm experiment is perhaps the longest-running, best-controlled, and most well-known artificial selection study bearing on the evolution of mammalian behavior. As such, it enables a uniquely powerful window on the neural mechanisms governing behavioral adaptation. This study used high-resolution MRI imaging to examine the brains of these foxes. We found that selection on social behavior has altered the anatomy of distributed gray matter networks which included, among other regions, prefrontal cortex, hippocampus, amygdala, caudate, nucleus accumbens, cerebellum, and hypothalamus. This agrees generally with past findings in these foxes^19–23^ and on the neural correlates of wolf-to-dog domestication^8–10,19–21,30–33^. Although some of our prefrontal results are located on the dorsolateral surface of the brain (i.e., the prorean gyrus), this region is likely not homologous to the granular dorsolateral prefrontal cortex of humans and macaque monkeys, as that region is thought to be unique to primates^34,35^. Rather, the connectivity and cytoarchitecture of carnivore prefrontal cortex^11,36–41^ are more similar to that of the dysgranular and agranular portions of primate orbitofrontal and ventromedial prefrontal cortex, regions which function to integrate external multisensory and internal visceromotor information with limbic reward and threat signals in order to compare the value of potential behavioral choices^42,43^. Putatively, this circuitry has been altered in tame foxes to bias behavioral decisions toward the reward value of social contact with humans, and in aggressive foxes toward the opposite. This interpretation fits generally with conceptualizations of social approach-avoidance processing in humans and other species^44^. It also fits with prior work suggesting that similar genetic changes might underlie shifts toward both aggression and tameness in these foxes, resulting in similar physiological and morphological changes^45^.

However, the current study also produced some findings which were unexpected and suggest revision of existing thinking about domestication. One of these was that total gray matter volume in both the tame strain and the aggressive strain is *increased* compared to the conventional strain. These findings are in contrast to a number of prior studies which have reported that domestication reduces brain size in diverse species including Atlantic cod^46^, guppies^47^, rainbow trout^48^, mallard ducks^49^, rats, mice, gerbils, guinea pigs, rabbits, pigs, sheep, llamas, horses, ferrets, cats, and dogs (reviewed in ^5^). Brain size should be interpreted in the context of body size, given that the two covary. While body size measurements were not available for the foxes in our study, Huang et al.^23^ report a 15.9% reduction in body weight in tame compared to conventional foxes, suggesting that if anything, differences in brain:body size ratios are more pronounced than the brain measurements alone would indicate. What might cause this increase in brain size? One possible explanation might involve neuroplastic changes resulting from differential lifetime experience. The link between naturalistic, enriched environments and larger, more complex brains has been noted for decades^50,51^. Tame, aggressive, and conventional foxes are all housed in identical conditions and undergo identical treatment. Nonetheless, tame foxes, because of their innate predisposition toward prosocial interaction with humans, may effectively experience this environment to be more naturalistic and enriched.

However, this potential interpretation does not explain why the aggressive strain *also* shows increased gray matter volume relative to controls. Perhaps their constant drive to avoid human contact functions as a sort of enrichment, but we instead favor an alternative explanation. Both the tame and aggressive strains have been subject to intense, sustained selection on behavior, while the conventional strain undergoes no such intentional selection. Thus, it is possible that fast evolution of behavior, at least initially, may generally proceed via increases in gray matter. Two mechanisms of brain evolution might explain this effect. First, because of developmental linkages, selection pressure for increased size in one region may “drag along” enlargement in others^52^. Second, due to the complex interdependencies between brain systems, adaptive solutions which modify one existing system may stand a high likelihood of producing deleterious effects in others unless additional compensatory adaptations also occur^53^. In the wild – and perhaps also in later stages of domestication, as animals are continually selected for increased economic efficiency – there would be constant pressure against these “easy, wasteful” solutions because of the metabolic and life-history constraints associated with increased brain size^54^. In the context of the fox-farm experiment, however, these constraints may have been lessened or removed. There are multiple potential cellular-level causes for increased gray matter, including increased neuron count, increased neuron size, and/or dendritic changes such as increased arborization or spine density^55,56^. Future histological research, including single cell transcriptomics and histological work, will be required to differentiate among these possibilities.

Notably, our results do agree with a recent skeletal morphology study which did not find a significant difference in endocranial volume between tame, aggressive, and conventional foxes (Kistner et al, in press). The foxes that became the progenitor population for Belyaev’s study had existed in a farmed state in Russia for about 50 years^57,58^; this original Russian farmed population was itself drawn largely from Eastern Canadian foxes^59^ bred at fur farms on Prince Edward Island^60^. Kistner et al. (in press) compared endocranial volumes of modern Russian fox farm skulls to skulls from wild foxes which had been collected in Canada east of Quebec between 1984-1952, with 70% collected between 1894-1900. This revealed that endocranial volume in the modern, conventional Russian farm-raised foxes was significantly reduced compared to the archived Canadian wild skulls. Together with these findings, the current study hints that the brain volume reduction associated with domestication might occur relatively quickly, prior to the onset of intentional selection on behavior. Further research is required to establish whether this is true.

A second surprising result from this study was that both tame and aggressive foxes showed enlargement in substantially overlapping gray matter regions, including prefrontal cortex, amygdala, hippocampus, and cerebellum (**Figure 2E**). In addition to these volumetric effects, we also observed strainwise differences in the degree of morphological covariation across distributed, multi-region networks. These latter measurements revealed links with individual variation in behavior, including in brain regions that did not show volumetric differences, notably, the hypothalamus. Additionally, Network 3 consisted primarily of the hypothalamus and prefrontal cortex, two regions strongly implicated in both fox and dog domestication^8–10,19–21,30–33^. In this network, factor loadings did not differentiate the tame from aggressive strains; rather, the selectively-bred strains together were differentiated from the conventional strain (**Figure 3C**). Taken together, these results indicate that selection for opposite behavioral responses (docility versus aggression) can produce similar adaptations in the brain. This has important implications for attempts to evaluate the hypothesis that humans are ourselves self-domesticated^61^, given that similar neuroanatomical patterns of change could now be interpreted to support either selection for increased *or* decreased aggression in our lineage. This calls for additional research on the neural correlates of evolved differences in both docility and aggression at the cellular and genomic levels.

The Russian fox-farm experiment represents a uniquely well-controlled opportunity to study the effects of specific, sustained selection on behavior. Thus, apart from questions of domestication, an additional implication of these results concerns brain evolution on a more general level. We found that intense selection on behavior can produce gross changes in distributed brain morphology extremely rapidly – within the span of well under a hundred generations. This suggests that the brains of other animals on this planet, including *Homo sapiens*, may have undergone similarly precipitous morphological shifts any time steep selection on behavior was experienced.

## Online Methods

### Brain specimens

In the current study, we examined the brains of 10 tame, 10 conventional, and 10 avoidant/aggressive foxes. Right hemispheres were preserved for gene expression studies; we report analyses in left hemispheres here. All foxes were male, sexually naïve, and approximately 1.5 years old. Brains were formalin-fixed.

### Neuroimaging data acquisition

For imaging, brains were placed in a waterproof plastic container which was packed with polyethylene beads for stabilization. The container was then pumped full of Fluorinert^TM^ FC-770 (3M^TM^). Fluorinert is a fluorocarbon; it is analogous to a hydrocarbon but with fluorine taking the place of hydrogen. It thus produces no signal in (typical) MRI imaging, which is tuned to the resonant frequency of hydrogen nuclei. Images were acquired on a 9.4 T/20 cm horizontal bore Bruker magnet, interfaced to an Avance console, with Paravision 5.1 software (Bruker, Billerica, MA, USA). A 7.2 cm diameter volume r.f. coil was used for transmission and reception with a RARE T2 sequence (2 averages, 13 ms echo time, 2500 ms repetition time, rare factor 8). Image resolution was 300 μm^3^ with a matrix size of 256 x 100 x 88.

### Image analysis

Image pre-processing was accomplished using the FSL software package^62–64^. Images underwent bias correction and segmentation into white matter and gray matter using *FAST*^65^. In order to provide a common spatial framework for morphometric analysis, an unbiased nonlinear template was built from the 10 conventional foxes’ T2-weighted images using the ANTS software package^66^. This template represents the group average morphology across the conventional fox brain specimens. All subject’s T2-weighted image were nonlinearly aligned to this template. We computed the Jacobian determinant image of each deformation field; this represents a spatial map of where and how much each individual subject’s scan had to deform in order to come into alignment with the template. The Jacobian determinant images were then masked with the gray matter segmentation images in order to produce representations of each subject’s gray matter deviation from the template. These images became the input for statistical morphometric analysis.

Two independent, complementary morphometric analysis approaches were used. Voxel based morphometry (VBM^67^) is an inherently hypothesis-based approach that carries out general linear modeling at each voxel in the image to determine whether morphology is significantly related to explanatory variables (such as group). VBM analysis was accomplished using FSL’s *randomise* tool for voxelwise Monte Carlo permutation testing of general linear models, which permutes explanatory variables across cases in order to build up a null distribution, and then tests whether observed associations to explanatory variables significantly differ from this random null distribution^68^. On the other hand, source-based morphometry (SBM^29^) is a data-driven, model-free approach that identifies patterns of significant morphological correlation across subjects. In other words, it determines which regions of the brain significantly covary with each other across the entire dataset, while remaining agnostic to putative differences across subjects (such as group differences). Post-hoc tests can then be used to determine whether these networks show significant associations with variables of interest. This was accomplished using the GIFT SBM toolbox for Matlab (^29^; http://mialab.mrn.org/software/gift/index.html). Note that although this approach identifies networks of structurally covarying regions, it does not assess white matter connections linking those regions.

### Assignment of fox behavioral phenotypes

Fox behavior was tested at 5.5–6 months of age in the standard test described in Kukekova et al.^24^. Fox behavior during the test was scored from the video records for 98 recordable observations (**Table S1**). The matrix including scores for 1003 foxes from Kukekova et al.^16^ and scores for 30 foxes used in this study were subjected to Principal Component analysis in R using function: *prcomp*.

## Acknowledgments

EEH, TPM, DAG, and neuroimaging scan costs were supported by NSF-IOS #1457291. AVK, sample collection, and behavioral data analysis were supported by NIH grant GM120782. We are grateful to Jaekeun Park and Orion Keifer for assistance with scan acquisition and to Olivia Zarella and Jeromy Dooyema for assistance with sample preparation. We thank Yury E. Herbeck and Anastasiya V. Kharlamova for insightful discussions and help in preparation of the experiments at the experimental farm of the Institute of Cytology and Genetics (ICG) in Novosibirsk, Russia. We are grateful to Irina V. Pivovarova, Anastasiya V. Vladimiriva, Tatyana I. Semenova, Eugene A. Martinov, and all the animal keepers at the ICG experimental farm for research assistance.

