## Supplementary material for "Neuromorphological changes following selection for tameness and aggression in the Russian fox-farm experiment": Table S1

**Table S1.** Trait loadings for principle components analysis of behavior.

| Trait | Trait description | PC1 | PC2 | PC3 |
|---|---|---|---|---|
| C37 | Aggressive sounds | -0.1476673 | 0.2057388 | 0.05690569 |
| A52 | Did not come to zone 2a | -0.1430533 | -0.0512895 | -0.0320838 |
| C34 | Follows the hand (aggr.) | -0.1402881 | 0.23863637 | 0.07126363 |
| C31 | Attack alert | -0.1399264 | 0.2314992 | 0.06913457 |
| C32 | Pinned ears (aggr.) | -0.133127 | 0.1853784 | 0.07517754 |
| D31 | Not on the floor of zone 2 | -0.1304852 | -0.0279166 | -0.1040545 |
| B12 | Not on the floor of zone 2 | -0.1286657 | -0.1034625 | 0.15847715 |
| A25 | Spend in zone 3-4-5-6 at least 40'' | -0.1145566 | -0.1145345 | -0.1041976 |
| C36 | Triangle ears directed back (aggr.) | -0.1084682 | 0.1639514 | 0.06852387 |
| B25 | Pinned ears (aggr.) | -0.1030547 | 0.12176326 | -0.0338697 |
| B29 | Spend in zone 3-4-5-6 at least 40'' | -0.0979483 | -0.1150948 | 0.18041277 |
| C30 | Attack | -0.0811832 | 0.20157786 | 0.02742591 |
| C33 | Trying to bite | -0.0787731 | 0.18749222 | 0.09395528 |
| B30 | Spend in zone 5-6 at least 40'' | -0.0763434 | -0.1219152 | -0.040511 |
| B42 | Keeping same posture and place for at least 40" | -0.0746656 | -0.0280051 | -0.1063443 |
| B13 | Not on the floor of zone 2a | -0.0730084 | -0.0776614 | 0.14535304 |
| B2 | Immediately moved back to zone 5 or 3-5-6 | -0.0667175 | -0.1315815 | 0.16132763 |
| A23 | Moving back for at least one zone during first 15" | -0.0635934 | -0.0798363 | 0.04627051 |
| A40 | Keep same posture and place at least for 40" | -0.0610294 | -0.0451188 | -0.0882064 |
| C38 | Animal is present only in zones 3-5-6 | -0.0603003 | -0.2143789 | 0.06897475 |
| D39 | Pinned ears (aggr.) | -0.0589739 | 0.07022877 | -0.0544917 |
| C4 | Spend more than 30" in zones 3-4-5-6 | -0.0486048 | -0.1129164 | 0.1191827 |
| C7 | First time can touch a fox in zones 5-6 | -0.0442354 | -0.2005908 | 0.16847207 |
| A31 | Lie in any zone longer than 30" | -0.0413531 | -0.0473826 | -0.0526588 |
| C3 | Animal is in zones 3-4-5-6 in the beginning of stepC | -0.0359193 | -0.0807357 | 0.10769476 |
| C55 | Leaning on side or back walls in zones 5-6 | -0.0357765 | -0.0621933 | 0.09883121 |
| A32 | Lie in any zone a whole minute | -0.0322557 | -0.0455808 | -0.0514736 |
| C35 | Narrow ears directed back | -0.0134525 | -0.1674438 | 0.07190769 |
| B37 | Stays on back or side walls in zones 5-6 | -0.0067197 | -0.0593972 | 0.07711811 |
| B48 | Ears are vertical | -0.0051633 | -0.0905863 | 0.21988206 |
| B14 | Sniffing floor/air | 0.00665074 | -0.0271967 | 0.12153536 |
| A48 | Sit for at least 20" | 0.00809613 | -0.029441 | -0.0223952 |
| C50 | Tail is up for at least for 3" | 0.01117132 | 0.03423211 | 0.11681401 |
| B9 | Sniffing hand from small distance | 0.01562686 | -0.0104944 | 0.12987889 |
| A34 | Changed place at least once | 0.02762113 | 0.01619735 | 0.0878816 |
| A38 | Did at least one full circle | 0.03153232 | -0.0493083 | 0.12161341 |

| | | | | |
|---|---|---|---|---|
| B8 | Come to the hand after 10" | 0.03582494 | -0.0003365 | 0.07587808 |
| A36 | Changed place at least 2-4 times | 0.03677646 | 0.01294125 | 0.10609875 |
| D13 | Groomming | 0.03813346 | -0.0131906 | 0.00017673 |
| B39 | Changed place at least 2-4 times | 0.0453691 | 0.00602985 | 0.0943546 |
| C2 | Animal is in zones 1-2-3-4 in the beginning of stepC | 0.04645646 | 0.10165381 | -0.0895807 |
| A47 | Tail is up for at least 3" | 0.05655872 | 0.00542154 | 0.05048287 |
| B47 | Tail is up for at least 3" | 0.05926645 | 0.00639307 | 0.08498355 |
| B32 | Moving on "short legs" | 0.06011035 | 0.0280033 | -0.0825452 |
| C39 | Moved forward for at least one zone during the step | 0.06502536 | 0.13752432 | 0.04538581 |
| C204 | Tame sounds (combined) | 0.06680337 | 0.00121025 | -0.0599009 |
| D33 | Tail is up for at least 3" | 0.0674776 | -0.0014009 | 0.01885399 |
| B19 | Loud breathing | 0.06814984 | 0.0352152 | -0.124301 |
| B3 | Touch hand for at least 40" | 0.06946151 | 0.04423451 | -0.1249051 |
| D25 | Moving forward for at least one zone during first 15'' | 0.07393203 | -0.0237983 | 0.10749179 |
| B1 | Animal is in zone 2 in the beginning | 0.07482935 | 0.08558415 | -0.1237285 |
| C25 | Tail wagging | 0.07724832 | -0.0012107 | -0.0996494 |
| C6 | First time can touch a fox in zones 3-4 | 0.0797726 | -0.0387303 | 0.01207579 |
| C18 | Hold hand | 0.08141347 | -0.0092092 | -0.061069 |
| D29 | Did at least one full circle | 0.084554 | -0.0796958 | 0.09728685 |
| A37 | Changed place at least 5 times | 0.08626927 | 0.01591251 | 0.17384956 |
| C17 | Rolls on the side, ask to touch belly | 0.08742998 | 7.45E-05 | -0.0816944 |
| A7 | Touch door by foot or scratch | 0.08991397 | 0.0532481 | -0.0091076 |
| A29 | Came to zone 1-2 | 0.09018569 | 0.0465907 | 0.15163891 |
| B40 | Changed place at least 5 times | 0.09072512 | -0.0165086 | 0.1744323 |
| C13 | Allows to touch back part of the back | 0.09318948 | -0.2048051 | 0.00241271 |
| D24 | Comes to zones 1-2 | 0.0959737 | 0.01736351 | 0.12844973 |
| A2 | Tail wagging | 0.10156213 | 0.02488945 | -0.0929812 |
| D28 | Changes place at least 5 times | 0.10330463 | -0.0334257 | 0.11796783 |
| D6 | Touch door by foot or scratch | 0.10455025 | 0.01406251 | -0.0169301 |
| B31 | Spend in zone 1-2 at least 40'' | 0.1063003 | 0.09768157 | -0.1496475 |
| B21 | Tame ears | 0.10675677 | 0.03203084 | -0.1522048 |
| B20 | Tail wagging | 0.10715258 | 0.03589231 | -0.1538206 |
| D3 | Tail wagging | 0.1081683 | -0.0032861 | -0.1271162 |
| A24 | Spend in zone 1-2-3-4 at least 40'' | 0.10833383 | 0.10252039 | 0.15880762 |
| A22 | Moving forward for at least one zone during first 15" | 0.11001501 | 0.0641225 | 0.15359587 |
| C24 | Loud breathing | 0.1132731 | -0.0010303 | -0.1092512 |
| A28 | Spend in zone 1-2 at least 10'' | 0.11350046 | 0.06669391 | 0.15984224 |
| A6 | Sniffing the door | 0.11575323 | 0.0628932 | 0.17880495 |
| C14 | Allows to touch back | 0.11793174 | -0.2310268 | -0.0218957 |

| | | | | |
|---|---|---|---|---|
| A9 | Lean on the right wall in zone 2 | 0.11959853 | 0.07496327 | 0.07927634 |
| D7 | Touch door by nose | 0.12007958 | 0.02927169 | 0.13297053 |
| D14 | Sit in zone 2 and looking on observer | 0.1224556 | 0.01267484 | -0.0627118 |
| A10 | Sit in zone 2 and looking on observer | 0.12628208 | 0.03313335 | -0.007816 |
| B28 | Spend in zone 1-2-3-4 at least 40'' | 0.12730388 | 0.13902538 | -0.1224083 |
| D17 | Spends in zone 1-2-3-4 at least 40" | 0.13379193 | 0.06232127 | 0.09512188 |
| D32 | Leaning on right wall in zone 2 | 0.13470139 | 0.00703826 | 0.02006886 |
| A8 | Lean on the door | 0.1363755 | 0.06708218 | 0.07616227 |
| A27 | Spend in zone 1-2 at least 40'' | 0.13917388 | 0.09237863 | 0.08274502 |
| C16 | Allows to touch head | 0.14020198 | -0.2408099 | -0.0473998 |
| A5 | Touch door with nose | 0.14037341 | 0.08793027 | 0.17366217 |
| C8 | Lie during a contact for at least 5" | 0.14106738 | -0.1907798 | -0.0817082 |
| B15 | Sniffing front door | 0.14260148 | 0.10017517 | 0.05436706 |
| C15 | Allows to touch nose | 0.14612813 | -0.2334574 | -0.0457441 |
| D1 | Come to the zone 2 during first 5" | 0.14915423 | 0.0313106 | 0.08782056 |
| C19 | Come to zone 2 in the end of stepC | 0.15108775 | 0.08114312 | -0.0033427 |
| D2 | Spends in zones 1-2 at least 30" | 0.15262612 | 0.03396873 | 0.0881678 |
| B7 | Trying to poke hand by nose | 0.15360996 | 0.06452159 | -0.0786887 |
| D4 | Leaning on door | 0.15681275 | -0.0184744 | 0.02205177 |
| C29 | Come to hand and sniffing in the end of stepC | 0.15705644 | 0.02431407 | -0.0027936 |
| B11 | Come to zone 1-2 | 0.15826159 | 0.1000979 | 0.00695365 |
| C12 | Tame ears | 0.1633044 | -0.1334769 | -0.0702261 |
| B10 | Come to hand and sniffing | 0.16338574 | 0.06024868 | -0.0512387 |
