## Supplementary material for "Neuromorphological changes following selection for tameness and aggression in the Russian fox-farm experiment": Table S2

**Table S2.** Clusters resulting from VBM analysis. Each component contains multiple clusters, indicated by *Cluster Index* . Voxel coordinates are for maximum value in cluster.

| | Cluster Index | Volume (mm3) | MAX X (vox) | MAX Y (vox) | MAX Z (vox) |
|---|---|---|---|---|---|
| **Tame > Aggressive** | **Cluster Index** | **Volume (mm3)** | **MAX X (vox)** | **MAX Y (vox)** | **MAX Z (vox)** |
| | 1 | 31.941 | 56 | 162 | 77 |
| | | | 55 | 160 | 79 |
| | | | 59 | 140 | 86 |
| | | | 62 | 132 | 82 |
| | | | 55 | 150 | 86 |
| **Tame < Aggressive** | **Cluster Index** | **Volume (mm3)** | **MAX X (vox)** | **MAX Y (vox)** | **MAX Z (vox)** |
| | 2 | 198.288 | 62 | 189 | 50 |
| | | | 70 | 186 | 52 |
| | | | 61 | 196 | 52 |
| | | | 55 | 193 | 52 |
| | | | 59 | 175 | 59 |
| | | | 72 | 180 | 72 |
| | 1 | | 79 | 139 | 76 |
| | | | 74 | 141 | 73 |
| | | | 68 | 154 | 63 |
| **Tame > Conventional** | **Cluster Index** | **Volume (mm3)** | **MAX X (vox)** | **MAX Y (vox)** | **MAX Z (vox)** |
| | 3 | 1095.957 | 32 | 222 | 69 |
| | | | 33 | 223 | 67 |
| | | | 36 | 222 | 78 |
| | | | 40 | 226 | 56 |
| | | | 46 | 207 | 64 |
| | | | 56 | 210 | 86 |
| | 2 | 303.183 | 34 | 98 | 59 |
| | | | 32 | 85 | 77 |
| | | | 35 | 73 | 79 |
| | | | 35 | 86 | 62 |
| | | | 48 | 100 | 47 |
| | | | 42 | 96 | 46 |
| | 1 | 185.22 | 36 | 129 | 27 |
| | | | 36 | 151 | 18 |
| | | | 54 | 122 | 59 |
| | | | 38 | 141 | 33 |
| | | | 39 | 145 | 27 |
| | | | 44 | 133 | 38 |
| **Tame < Conventional** | **(none)** | | | | |
| **Aggressive > Conventional** | **Cluster Index** | **Volume (mm3)** | **MAX X (vox)** | **MAX Y (vox)** | **MAX Z (vox)** |
| | 11 | 2309.283 | 37 | 222 | 79 |
| | | | 63 | 190 | 49 |
| | | | 31 | 223 | 70 |
| | | | 61 | 198 | 51 |
| | | | 56 | 202 | 58 |
| | | | 29 | 217 | 93 |

| | | | | | |
|---|---|---|---|---|---|
| | 10 | 440.505 | 42 | 130 | 130 |
| | | | 78 | 123 | 104 |
| | | | 41 | 121 | 130 |
| | | | 77 | 96 | 90 |
| | | | 42 | 140 | 129 |
| | | | 42 | 143 | 128 |
| | 9 | 104.652 | 48 | 149 | 17 |
| | | | 49 | 141 | 17 |
| | | | 54 | 149 | 18 |
| | | | 52 | 145 | 27 |
| | | | 39 | 148 | 17 |
| | | | 35 | 129 | 26 |
| | 8 | 93.717 | 54 | 125 | 51 |
| | | | 61 | 122 | 51 |
| | | | 46 | 141 | 37 |
| | 7 | 52.083 | 36 | 90 | 70 |
| | | | 44 | 86 | 69 |
| | | | 46 | 77 | 75 |
| | | | 53 | 79 | 70 |
| | 6 | 22.896 | 81 | 132 | 75 |
| | | | 80 | 139 | 75 |
| | | | 78 | 142 | 65 |
| | 5 | 9.855 | 52 | 107 | 70 |
| | | | 46 | 103 | 63 |
| | 4 | 3.402 | 26 | 163 | 58 |
| | | | 30 | 157 | 62 |
| | 3 | 3.348 | 33 | 74 | 82 |
| | 2 | 0.567 | 64 | 145 | 59 |
| | 1 | 0.486 | 42 | 206 | 30 |
| **Aggressive < Conventional** | **(none)** | | | | |
