## Supplementary material for "Neuromorphological changes following selection for tameness and aggression in the Russian fox-farm experiment": Table S3

**Table S3.** Clusters resulting from SBM analysis. Within each network, the negative components (shown as blue in Fig. 3) and positive components (red) are morphologically anti-correlated. Each component contains multiple clusters, indicated by *Cluster Index* . Voxel coordinates are for maximum value in cluster.

| **Network 1 (negative/blue)** | **Cluster Index** | **Volume (mm3)** | **MAX X (vox)** | **MAX Y (vox)** | **MAX Z (vox)** |
|---|---|---|---|---|---|
| | 18 | 137.997 | 14 | 114 | 120 |
| | | | 13 | 116 | 114 |
| | | | 13 | 95 | 122 |
| | | | 14 | 126 | 122 |
| | | | 14 | 150 | 123 |
| | 17 | 125.712 | 29 | 135 | 68 |
| | | | 25 | 145 | 63 |
| | | | 21 | 149 | 70 |
| | | | 19 | 160 | 61 |
| | | | 19 | 158 | 56 |
| | | | 30 | 152 | 72 |
| | 16 | 48.573 | 61 | 80 | 51 |
| | 15 | 39.312 | 52 | 218 | 53 |
| | | | 44 | 210 | 34 |
| | | | 52 | 204 | 44 |
| | 14 | 17.523 | 40 | 219 | 87 |
| | | | 32 | 208 | 97 |
| | | | 24 | 202 | 97 |
| | 13 | 8.451 | 61 | 213 | 70 |
| | 12 | 6.129 | 34 | 215 | 19 |
| | 11 | 5.265 | 18 | 87 | 48 |
| | 10 | 5.238 | 15 | 118 | 86 |
| | 9 | 5.184 | 60 | 86 | 68 |
| | 8 | 2.349 | 41 | 122 | 50 |
| | 7 | 1.755 | 65 | 156 | 53 |
| | 6 | 1.188 | 66 | 195 | 57 |
| | | | 69 | 192 | 61 |
| | | | 70 | 190 | 62 |
| | 5 | 1.053 | 26 | 209 | 85 |
| | 4 | 0.837 | 54 | 155 | 51 |
| | 3 | 0.567 | 33 | 233 | 30 |
| | 2 | 0.459 | 18 | 107 | 56 |
| | 1 | 0.432 | 78 | 99 | 69 |

| **Network 1 (positive/red)** | **Cluster Index** | **Volume (mm3)** | **MAX X (vox)** | **MAX Y (vox)** | **MAX Z (vox)** |
|---|---|---|---|---|---|
| | 14 | 383.94 | 24 | 51 | 89 |
| | | | 30 | 55 | 83 |
| | | | 32 | 53 | 70 |
| | | | 34 | 56 | 75 |
| | | | 26 | 49 | 68 |
| | | | 17 | 102 | 78 |

| | | | | | |
|---|---|---|---|---|---|
| | 13 | 94.338 | 40 | 234 | 57 |
| | | | 41 | 229 | 49 |
| | | | 35 | 225 | 73 |
| | | | 33 | 238 | 59 |
| | | | 30 | 234 | 69 |
| | | | 37 | 214 | 44 |
| | 12 | 41.499 | 43 | 73 | 42 |
| | | | 47 | 61 | 55 |
| | 11 | 24.516 | 59 | 105 | 60 |
| | | | 66 | 108 | 54 |
| | 10 | 17.604 | 48 | 99 | 44 |
| | 9 | 16.659 | 23 | 118 | 103 |
| | | | 30 | 113 | 100 |
| | | | 40 | 107 | 100 |
| | 8 | 6.831 | 49 | 201 | 94 |
| | 7 | 4.617 | 28 | 178 | 36 |
| | 6 | 3.996 | 50 | 101 | 77 |
| | 5 | 3.402 | 36 | 186 | 102 |
| | 4 | 2.052 | 57 | 134 | 51 |
| | 3 | 0.864 | 28 | 142 | 57 |
| | 2 | 0.162 | 26 | 157 | 45 |
| | 1 | 0.027 | 26 | 151 | 40 |

| **Network 2 (negative/blue)** | **Cluster Index** | **Volume (mm3)** | **MAX X (vox)** | **MAX Y (vox)** | **MAX Z (vox)** |
|---|---|---|---|---|---|
| | 9 | 153.954 | 21 | 55 | 95 |
| | | | 17 | 88 | 93 |
| | | | 16 | 97 | 85 |
| | 8 | 40.635 | 24 | 158 | 38 |
| | 7 | 39.501 | 71 | 173 | 88 |
| | | | 61 | 156 | 109 |
| | | | 62 | 158 | 108 |
| | | | 68 | 168 | 96 |
| | | | 65 | 133 | 111 |
| | | | 66 | 131 | 110 |
| | 6 | 33.966 | 73 | 161 | 83 |
| | | | 74 | 154 | 82 |
| | 5 | 25.596 | 73 | 97 | 54 |
| | | | 69 | 85 | 64 |
| | | | 67 | 82 | 70 |
| | 4 | 12.15 | 33 | 232 | 76 |
| | | | 33 | 237 | 68 |
| | | | 32 | 236 | 70 |
| | 3 | 9.18 | 23 | 44 | 70 |
| | 2 | 0.378 | 26 | 99 | 100 |
| | 1 | 0.054 | 56 | 112 | 40 |

| **Network 2 (positive/red)** | **Cluster Index** | **Volume (mm3)** | **MAX X (vox)** | **MAX Y (vox)** | **MAX Z (vox)** |
|---|---|---|---|---|---|

| | | | | | |
|---|---|---|---|---|---|
| | 11 | 528.201 | 55 | 76 | 49 |
| | | | 47 | 84 | 39 |
| | | | 51 | 98 | 41 |
| | | | 33 | 55 | 55 |
| | | | 43 | 95 | 54 |
| | | | 35 | 56 | 59 |
| | 10 | 48.168 | 18 | 90 | 50 |
| | | | 19 | 92 | 65 |
| | | | 23 | 90 | 74 |
| | 9 | 8.289 | 63 | 145 | 77 |
| | 8 | 7.128 | 33 | 75 | 74 |
| | 7 | 5.373 | 15 | 118 | 88 |
| | 6 | 1.89 | 66 | 135 | 82 |
| | 5 | 1.674 | 33 | 223 | 54 |
| | 4 | 0.945 | 18 | 114 | 106 |
| | 3 | 0.648 | 39 | 216 | 44 |
| | 2 | 0.027 | 44 | 218 | 66 |
| | 1 | 0.027 | 49 | 63 | 75 |

| **Network 3 (negative/blue)** | **Cluster Index** | **Volume (mm3)** | **MAX X (vox)** | **MAX Y (vox)** | **MAX Z (vox)** |
|---|---|---|---|---|---|
| | 4 | 4.428 | 17 | 90 | 53 |
| | 3 | 2.241 | 22 | 47 | 55 |
| | | | 23 | 43 | 58 |
| | 2 | 0.729 | 18 | 95 | 62 |
| | 1 | 0.324 | 19 | 89 | 74 |

| **Network 3 (positive/red)** | **Cluster Index** | **Volume (mm3)** | **MAX X (vox)** | **MAX Y (vox)** | **MAX Z (vox)** |
|---|---|---|---|---|---|
| | 4 | 898.344 | 41 | 225 | 71 |
| | | | 50 | 217 | 82 |
| | | | 58 | 212 | 69 |
| | | | 32 | 228 | 54 |
| | | | 41 | 229 | 46 |
| | | | 59 | 203 | 86 |
| | 3 | 11.205 | 22 | 139 | 36 |
| | 2 | 3.159 | 27 | 156 | 39 |
| | 1 | 0.513 | 37 | 103 | 124 |

| **Network 4 (negative/blue)** | **Cluster Index** | **Volume (mm3)** | **MAX X (vox)** | **MAX Y (vox)** | **MAX Z (vox)** |
|---|---|---|---|---|---|
| | 14 | 14.31 | 46 | 80 | 63 |
| | | | 41 | 81 | 69 |
| | 13 | 12.285 | 72 | 152 | 92 |
| | | | 74 | 163 | 83 |
| | 12 | 9.531 | 53 | 79 | 43 |
| | | | 52 | 84 | 49 |
| | 11 | 9.018 | 76 | 121 | 55 |
| | 10 | 4.239 | 49 | 98 | 64 |
| | 9 | 2.106 | 32 | 238 | 62 |

| | Cluster Index | Volume (mm3) | MAX X (vox) | MAX Y (vox) | MAX Z (vox) |
|---|---|---|---|---|---|
| | 8 | 1.377 | 45 | 64 | 66 |
| | 7 | 1.188 | 32 | 233 | 75 |
| | 6 | 0.918 | 39 | 107 | 45 |
| | 5 | 0.783 | 60 | 87 | 67 |
| | 4 | 0.621 | 50 | 100 | 82 |
| | 3 | 0.216 | 28 | 125 | 74 |
| | 2 | 0.081 | 54 | 74 | 52 |
| | 1 | 0.027 | 74 | 107 | 53 |

| **Network 4 (positive/red)** | **Cluster Index** | **Volume (mm3)** | **MAX X (vox)** | **MAX Y (vox)** | **MAX Z (vox)** |
|---|---|---|---|---|---|
| | 12 | 452.682 | 24 | 156 | 38 |
| | | | 21 | 139 | 36 |
| | | | 26 | 130 | 41 |
| | | | 26 | 143 | 59 |
| | | | 21 | 127 | 35 |
| | | | 21 | 152 | 52 |
| | 11 | 184.788 | 17 | 97 | 53 |
| | | | 17 | 114 | 71 |
| | | | 20 | 90 | 74 |
| | | | 16 | 126 | 60 |
| | | | 15 | 112 | 65 |
| | | | 16 | 111 | 55 |
| | 10 | 98.685 | 26 | 46 | 68 |
| | | | 25 | 46 | 51 |
| | | | 24 | 46 | 49 |
| | 9 | 26.109 | 27 | 117 | 130 |
| | | | 26 | 112 | 129 |
| | | | 28 | 107 | 128 |
| | | | 27 | 133 | 131 |
| | | | 27 | 141 | 130 |
| | 8 | 10.773 | 24 | 65 | 78 |
| | 7 | 10.746 | 23 | 67 | 50 |
| | 6 | 1.539 | 36 | 94 | 122 |
| | 5 | 0.486 | 32 | 135 | 92 |
| | 4 | 0.27 | 21 | 63 | 38 |
| | 3 | 0.162 | 56 | 215 | 63 |
| | 2 | 0.081 | 21 | 66 | 94 |
| | 1 | 0.027 | 35 | 134 | 85 |
